# Conserved genes with variable expression and function: *Vasa*, *Piwi*, and *Dnmt1* in *Oncopeltus fasciatus* gametogenesis

**DOI:** 10.64898/2026.09.11.751002

**Authors:** Patricia J. Moore, Emily A. Shelby, Elizabeth C. McKinney, Christopher B. Cunningham, Allen J. Moore

**Affiliations:** Department of Entomology, University of Georgia, Athens, GA 30602

**Keywords:** oogenesis, plasticity of germ cell gene set, spermatogenesis, reproductive genes

## Abstract

The production of gametes is one of the universal processes of life and given millennia of evolution many of the core genes involved are expected to be highly refined and resistant to change. Here we examine the pattern of expression and functionally characterize three genes widely involved in gametogenesis: *Vasa*, *Piwi*, and *Dnmt1*. We examine their cellular location during both oogenesis and spermatogenesis in *Oncopeltus fasciatus*, a hemipteran insect, using fluorescent *in situ* hybridization chain reaction. In females, expression of Piwi was restricted to the trophocytes, but *Vasa* and *Dnmt1* were expressed in developing ooctyes as well as the trophoctyes. In males, *Piwi* was expressed in the testis germline stem cells (GSCs), but *Vasa* and *Dnmt1* expression was absent from these cells. Additionally, *Piwi* and *Vasa* were expressed in secondary spermatogonia and spermatocytes while *Dnmt1* was predominantly expressed in the primary spermatogonia. We also functionally characterized their necessity for gametogenesis after RNAi-mediated gene expression knockdown. *Vasa* was not required for oogenesis but was required during spermatogenesis. *Piwi* and *Dnmt1* were required for both oogenesis and spermatogenesis. These results were in contrast to their requirement for these processes from other organisms, which highlight the unexpected variation found in the processes and suggest that conservation depends on the level at which genes are examined: sequence, expression, or phenotypic and biochemical function. This suggests there is a more nuanced evolutionary story of conserved, yet plastic, gametogenic gene set in Metazoa that deserves further study across more organisms.

**Summary statement:** Analysis of the expression patterns and function of *Vasa*, *Piwi*, and *Dnmt1* during gametogenesis in *Oncopeltus fasciatus* revealed that their roles varied when compared to other species, underscoring the importance of examining conservation of reproductive genes at multiple biological levels, in organisms with diverse reproductive ecology, and among the sexes.

## Introduction

The production of gametes is a foundational component of fitness and thus we predict that gametogenesis will be resistant to evolutionary change (Hirsh and Fraser 2001). This prediction is borne out by the observation that a core set of genes involved in the germ line and gametogenesis are highly conserved across animals (Ewen Campen et al. 2010, Fierro-Constaín et al. 2017, Brattig-Correia 2024). For example, both *Vasa* and *P-element induced wimpy testis* (*Piwi*) are highly conserved genes involved in gametogenesis, expressed in germ cells widely across the Metazoa (e.g. Schwager et al. 2015 and Chen et al. 2025). However, when the specific functional pathways of conserved genes is analyzed, differences among species are apparent, even within taxonomic groups. For example, while *Vasa* is involved in germline specification in both *Drosophila melanogaster* and the pea aphid *Acyrthosiphon pisum*, *Vasa* relies on *Oskar* in *D. melanogaster* but acts through an *Oskar*-independent mechanism in *A. pisum* (Lin and Chang 2025). Recently another gene, *DNA methyltransferase 1* (*Dnmt1*), has been shown to exhibit a similar expression pattern to *Piwi* and *Vasa* during insect gametogenesis, both in playing a required role but with variable function in reproduction across the insects (Zwier et al. 2012, Kay et al. 2018, Schulz et al. 2018, Bewick et al. 2019, Amukamara et al. 2020, Washington et al. 2021, Shelby et al. 2023). Thus, it appears that the evolutionary pattern of these genes is not the product of simple static conservation. This highlights the importance of defining at what level genes are conserved: DNA sequence, expression patterns, and function at the phenotypic and biochemical level.

Interpreting variation in these critical conserved gametogenesis gene sets and using it to better understand the evolution of reproduction is difficult due to incomplete data between species. For some species there is expression data, but no functional characterization. For other species, there might be functional characterization in oogenesis, but not spermatogenesis. Thus, discussing broad patterns of conserved presence yet variable gene function in gametogenesis is hampered. One way to address this is to begin to define their function within species with diverse reproductive physiologies and ecologies across the sexes. We designed this study to contribute to this effort by examining in detail the expression patterns and function of *Vasa*, *Piwi*, and *Dnmt1* in gametogenesis in the hemipteran *Oncopeltus fasciatus*, an insect commonly used as a model for insect development.

*Vasa* and *Piwi* are broadly recognized as two core members of the germline gene set (Ewen-Campen et al. 2010, Fierro-Constaín et al. 2017, Piccinni & Milani 2023, Chen et al. 2025, Kao et al. 2026) and both *Vasa* and *Piwi* are expressed in the germ cells of insects (Santos et al. 2023, Chen et al. 2025). *Vasa* was first identified in *D. melanogaster* where it is required for oogenesis (reviewed in Adeshev et al. 2023). However, the requirement for *Vasa* in gametogenesis is not universal even in the few cases where it has been functionally characterized within other arthropods. *Vasa* is required for oogenesis in spiders (Schwager et al. 2015) and sea lice (Bustos et al. 2023), but not in crickets (Ewen Campen 2013a) or *O. fasciatus* (Ewen Campen et al. 2013b). Similarly, *Vasa* is not required for spermatogenesis in *D. melanogaster* (Lasko 2013) but the role of *Vasa* during *D. melanogaster* male fertility may be more complex than previously thought (Adashev et al. 2024). *Vasa* is required for maintaining spermatogonia in crickets (Ewen Campen et al. 2013a). In *O. fasciatus*, knockdown of *Vasa* expression with RNA interference (RNAi) results in structural changes in the testis tubule, but the impact on male fertility was not tested (Ewen Campen et al. 2013b). Similar to *Vasa*, the function of *Piwi* in gametogenesis has only been functionally characterized in a few species of arthropods. The role of *Piwi* in gametogenesis has mainly been studied in *D. melanogaster* (Santos et al. 2023) where in both males and females *Piwi* is required for maintaining the germline stem cell population (Juliano et al. 2011). In a spider, *Parasteatoda tepidariorum*, *Piwi* is required for egg laying, but the function in spermatogenesis was not tested (Schwager et al. 2015). The picture in hemimetabolous insects is more complex. In the cricket *Gryllus bimaculatus*, *Piwi* is required for spermatogenesis, but not oogenesis (Ewen-Campen et al. 2013a). However, in the true bug *Rhodnius prolixus*, RNAi knockdown of *Piwi* results in disrupted oogenesis (Brito et al. 2018). The effect on male fertility was not examined. In *O. fasciatus*, another true bug, *Piwi* is strongly expressed in the tropharium of the ovary, but was not functionally characterized (Ewen-Campen et al. 2013b).

*DNA methyltransferase 1* (*Dnmt1*) is a highly conserved gene responsible for maintaining CpG DNA methylation patterns after DNA replication (Lyko 2018, Schmitz et al. 2019). Within the insects, however, *Dnmt1* conservation is variable (Bewick et al. 2017, Provataris et al. 2018, Duncan et al. 2022, Engelhart et al. 2022). Both CpG DNA methylation and the methylation toolkit are variable across the insects (Bewick et al. 2017, Provartaris et al. 2018) and the evolutionary stability of *Dnmt1* is not always tied to DNA methylation (Bewick et al. 2017). *Dnmt1* is expressed in the gonads of many species of insect and is functionally required for gametogenesis in a number of species (Zhang et al. 2015, Bewick et al. 2019, Amukamara et al. 2020, Washington et al. 2020, Ivasyk et al. 2023, Shelby et al. 2023, Cunningham et al. 2024), including *Tribolium castaneum*, a beetle with undetectable levels of CpG methylation (Zemach et al. 2010, Schulz et al. 2018). Differences of CpG methylation are not strongly or universally associated with gene expression difference across insects (Gladstad et al., 2019; Oldroyd & Yagound, 2021; Duncan et al., 2022; Maleszka & Kucharski, 2022; Bogan & Yi, 2024), including in *O. fasciatus* (Bewick et al., 2019, Washington et al., 2020), despite some exceptions (Arsala et al., 2022; Bain et al., 2021). This has led us and others to the hypothesis that there is a pleiotropic function for DNMT1 during gametogenesis in insects that is independent of DNA methylation (Amukamara et al., 2020; Ivasyk et al., 2025). Thus, *Dnmt1* in the insects generally demonstrates similar expression pattern and requirement as a core gametogenesis gene, but it shows variation across the insect tree of life and beyond too.

Using *O. fasciatus*, we combined experiments to examine both the expression patterns and function, through gene expression reduction using RNAi, of *Vasa*, *Piwi*, and *Dnmt1*. While *Vasa* is known not to be required for oogenesis in *O. fasciatus* (Ewen-Campen et al. 2013b), we show that it is also required for spermatogenesis. We found *Piwi*, which had not been previously functionally analyzed in *O. fasciatus*, was required for both oogenesis and spermatogenesis. *Dnmt1* is required for both oogenesis and spermatogenesis (Bewick et al. 2019, Amukamara et al. 2020, Washington et al. 2021). We also documented the specific cell types expressing these three genes within the ovary and testis. One highlight from our results was that *Piwi* is expressed in the testis germline stem cells (GSCs), but both *Vasa* and *Dnmt1* expression was absent from these cells. Finally, we refined the phenotype of the RNAi knockdowns by examining the impact of knockdown on the progression of gametogenesis, providing a more detailed and complete analysis of the block to gamete formation.

The variation we see raises further questions around the meaning of “conserved genes.” We conclude our paper with a brief discussion of these results in light of our original prediction that genes tightly tied to fitness should be conserved, as any mutation that negatively affects gamete production will be selected against. What does it mean to be “conserved”? At what level do we expect conservation versus tolerance of variation? Finally, how do we reconcile the prediction of conservation with the observation that many reproductive genes evolve rapidly (Dapper & Wade 2020)?

## Materials and Methods

### Animal husbandry

We established *Oncopeltus fasciatus* colonies with insects purchased from Carolina Biological Supply (Burlington, NC). We maintained colonies with *ad libitum* deionized water, organic raw sunflower seeds (Food to Live, Brooklyn, NY), and absorbent cotton wool oviposition substrate for oviposition. All animals were under a 16:8 light:dark cycle at 26°C. We staged nymphs by the development of the wing pads and pigmentation (Chesebro et al. 2009) and collected and housed fourth instar nymphs in separate containers. We collected teneral adults daily and housed nymphs in single sexed boxes until needed for experiments.

### RNAi knockdowns

RNAi knockdown of our target genes, *Vasa*, *Piwi*, and *Dnmt1*, was performed using published methods (Amukamara et al. 2020, Washington et al. 2021). The *O. fasciatus* genome contains only a single copy of *Vasa* and *Piwi* genes (Ewen-Campen et al. 2013OF) and *Dnmt1* (Bewick et al. 2019). Sense and anti-sense RNA for each RNAi (primers found in Supplementary Materials Table S1) were transcribed together with an Ambion MEGAscript kit (ThermoFisher Sci, Waltham, MA) and allowed to anneal to form dsRNA. We used *eGFP* as an exogenous construct to control for general RNAi and injection effects. The concentration of dsRNA was adjusted to 150 ng/μL in injection buffer (5 mM KCl, 0.1 mM NaH2PO4). All individuals were injected in the abdomen with 3 μL of dsRNA solution (450 ng total dsRNA) using pulled glass capillary needles (Sutter Instrument Company micropipette puller model P-97, Novato, CA). Following injection, individuals were housed in petri dishes, provided with *ad libitum* deionized water and sunflower seeds for recovery until analysis.

### Effect of knockdown on fertility

To examine the impact of knockdown of *Piwi* and *Dnmt1* on oocyte maturation and fertility, females were injected with ds-RNA at 7-10-days post adult emergence (PAE). This isolated the effect of knockdown on ovary development (Amukamara et al. 2020) from the effect on oocyte maturation. We aimed for 12-15 biological replicates per treatment. This sample size is sufficient for determining variation in female fecundity, based on multiple previous studies from our laboratory (Bewick et al. 2019, Amukamara et al. 2020, Shelby et al. 2023, Cunningham et al. 2024.) Females injected with dsRNA at 7-10-day PAE were paired with a virgin 10-day PAE untreated male. Pairs were housed individually in a petri dish with *ad libitum* deionized water, sunflower seeds, and cotton wool. Eggs were collected every 3-4 days and monitored daily for hatching. Each clutch is recognizable as a cluster of eggs and the number of eggs in the clutch and the percent of eggs from each clutch that hatched was recorded. The data for the first three clutches of each treated female was collected. We analyzed the change in percent of eggs that hatched using repeated measures ANOVA in JMP Pro v19 with treatment as our factor. We did not examine *Vasa* and female fertility as Ewen-Campen et al. (2013b) have previously shown that *Vasa* knockdown does not affect female fertility.

Similar to the females, for fertility studies, males were injected as adults, at 5 days PAE. Because males emerge as adults with functional sperm stores (Economopolous and Gordon, 1971), we measured the loss of fertility of a male over time to assess whether or not males were able to replenish sperm stores through the production of sperm following RNAi knockdown. We aimed for 20-25 biological replicates per treatment. This sample size is sufficient for determining variation in male fertility, based on multiple previous studies from our laboratory (Washington et al. 2021, Cunningham et al. 2023.) Male fertility was assessed using the fecundity of sequential mating partners (Washington et al., 2021). Following treatment, males were paired with a 7–10-day PAE virgin female and allowed to mate for seven days. Following mating, we removed the female and placed a new a virgin 7–10-day PAE female for another seven days until all males had mated with a total of four females. After we removed each female, we housed the female individually in a petri dish with *ad libitum* water, food, and cotton wool to measure the hatching success of the eggs each male’s mate laid. We collected all eggs laid by each female across its lifespan. Thus, for each biological replicate, we collected data on fertilization rates from four females that had mated to the male in order (1^st^ through 4^th^ mate) and analyzed the change in male fertility over time using repeated measures ANOVA in JMP Pro v19 with treatment as our factor.

### Hybridization chain reaction (HCR)

We localized mRNA using hybridization chain reaction (HCR; Molecular Instruments, Los Angeles, CA) during gametogenesis to visualize expression patterns of our target genes. We dissected ovaries and testes from treated sexually mature adults and removed individual ovarioles and testis tubules from surrounding membranes (Amukamara et al. 2020, Washington et al. 2021). For analysis of ovary phenotype when the target genes were knocked down prior to sexual maturation, we injected fourth instar female nymphs. Female 4^th^ instar nymphs were isolated from the mass colonies. The stage of development was identified through wing pad development (Chesebro et al. 2009) and sexed. Nymphs were chilled for 15 minutes and then injected with 3 μL of dsRNA solution as described above. For analysis of testis phenotype, we injected 5-days PAE as we had found that earlier injections had severe phenotypes that masked specific effects (Washington et al. 2021.) Ovarioles or testis tubules from a minimum of 10 individuals from each treatment were pooled for every experiment. Staining was repeated multiple times as needed to obtain the required images.

We fixed the tissues overnight at 4°C in Carnoys fixative (6:3:1 ethanol: chloroform: acetic acid). After this, samples were washed three times and resuspended in pure ethanol and stored at −20°C until further processing. We rehydrated the samples over a series of methanol dilutions in 0.3% Tween-20 in PBS (PTw; 3:1, 1:1, then 1:3) immediately prior to hybridization, followed by three washes in PTw. We largely followed the protocol of Yoon et al. (2024) for HCR with a few modifications. Briefly, samples were prehybridized with HCR probe hybridization buffer (Molecular Instruments Inc) at 37°C for 30 minutes with slow rotation. Probe solution was prepared by adding 1 pmol of each sequence specific probe set to 100 μL probe hybridization buffer (Molecular Instruments, Los Angeles, CA) at 37° C. Pre-hybridization solution was replaced with the prepared probe solution and incubated overnight (16–18 h) at 37° C with low shaking. Excess probe was removed by four 15-min washes with HCR probe wash buffer (Molecular Instruments) at 37° C followed by two 5-min washes with 5X SSCT (5X SSC + 0.1% Tween 20) at RT. We then pre-amplified the samples in HCR amplification buffer for 30-min at RT. While the samples were in pre-amplification buffer, we prepared 3 pmol of hairpin 1 and 3 pmol of hairpin 2 (separately) for each of our probes by heating to 95° C for 90 seconds in a thermocycler and then allowing to cool, protected from light, for 30 minutes at RT. We removed the preamplification buffer from samples and replaced it with amplification buffer to which the appropriate hairpins had been added. We incubated the samples with the hairpins at RT in the dark overnight. The following day we removed excess hairpins by washing with 5X SSCT at RT with gentle shaking, protected from the light. The samples were washed two times 5-min with 5X SSCT alone followed by a 30-min wash with 5X SSCT with 1 mg/ml DAPI added as a nuclear counterstain. We finished with a 30-min and 5-min wash with 5X SSCT and before mounting the samples on glass slides with Prolong Diamond Antifade Mountant (ThermoFisher Instruments, Waltham, MA).

### Immunohistochemistry

Using the protocol in Amukamara et al. (2020), we dissected ovaries from 7–10-day PAE untreated or RNAi knockdown females into PBS. Individual ovarioles were separated from supporting tissues and fixed in PBT (PBS + 0.1% Triton X-100) for 25 minutes at RT. We removed the fixative and washed the samples three times with PBT. We incubated the samples with primary antibody against α-β-catenin [α-CTNNB1; Sigma Aldrich cat# HPA029159] diluted 1:1000 in PBT, followed by an Alexa Fluor 647 goat-anti-rabbit secondary antibody (ThermoFisher Instruments, Waltham, MA) and counterstained with 0.5 μg/mL DAPI in PBT.

We visualized both the HCR and antibody labelled samples with a Zeiss LSM 880 Confocal Microscope (Zeiss Oberkochen, Germany) at the UGA Biomedical Microscopy core.

### Gene expression analysis

We extracted RNA from ovaries and testes of treated, sexually mature adults with a Qiagen RNeasy mini plus kit with Qiazol (Qiagen, Venlo, The Netherlands) and synthesized complementary DNA (cDNA) using 500 ng RNA with qScript cDNA Super-Mix (Quanta Biosciences, Gaithersburg, MD) following each manufacturer’s instructions. We determined gene expression levels of each gene of interest using quantitative real-time PCR (qRT-PCR) using ten biological replicates per gene of interest. Primers for the genes analyzed are listed in Table 1. We used *actin* as our endogenous reference gene (Amukamara et al. 2020, Washington 2021). We used a Roche LightCycler 480 with the SYBR Green Master Mix (Roche Applied Science Indianapolis, IN). We ran all samples with three technical replicates. Primer efficiency calculations, genomic contamination testing, and endogenous reference gene selection were performed as described in Cunningham et al. 2014. We used the ΔΔCT method to compare levels of gene expression across the samples (Livak and Schmittgen 2001). Differences in gene expression levels were analyzed using ANOVA in JMP Pro v14. If there was a significant overall effect, we compared means using Dunnett’s Method with ds*eGFP* as our control.

## Results

### Knockdown of expression by RNAi was effective

We tested expression levels of all three target genes in the ovaries of sexually mature adult females that had been treated with dsRNA as fourth instar nymphs (Supplementary Materials Figure S1) and testes of adult males that had been treated with ds-RNA during sexual maturation (Supplementary Materials Figure S2). In each case, the gene targeted with the dsRNA had significantly reduced expression.

### Dnmt1 and Piwi affect female fertility through different mechanisms

Of our three target genes, Ewen-Campen et al. (2013) has already demonstrated that *Vasa* does not affect female fertility. In that study, the authors show that *Piwi* is expressed in ovarioles, particularly in the germarium, but they did not functionally analyze *Piwi* in that study. We therefore investigated the function of *Piwi*, along with *eGFP* and *Dnmt1* as controls, in oogenesis. Female fecundity following RNAi knockdown in sexually mature adults showed an interaction between clutch number and treatment (Figure 1; repeated measures ANOVA time*treatment *F* = 5.929, d.f. = 4, 70, p < 0.001) showing that the change over time depended on treatment. As has been previously observed, females treated with ds-*eGFP*, similarly to untreated females, have an egg hatch rate with a mean of 80% (Bewick et al. 2019) that persisted across time. Females treated with ds-*Dnmt1* have a very low hatch rate in the first clutch (17%), probably representing oocytes that had chorions at the time of treatment and therefore were refractory to the dsRNA, and the hatch rate dropped to 0% in subsequent clutches. Adult females treated with ds-*Piwi* showed a different pattern in fertility. The first two clutches had typical hatch rates as the ds-*eGFP* treated females (mean of 72%). However, the hatch rate was greatly reduced for the third clutch (41%).

**Figure 1.**
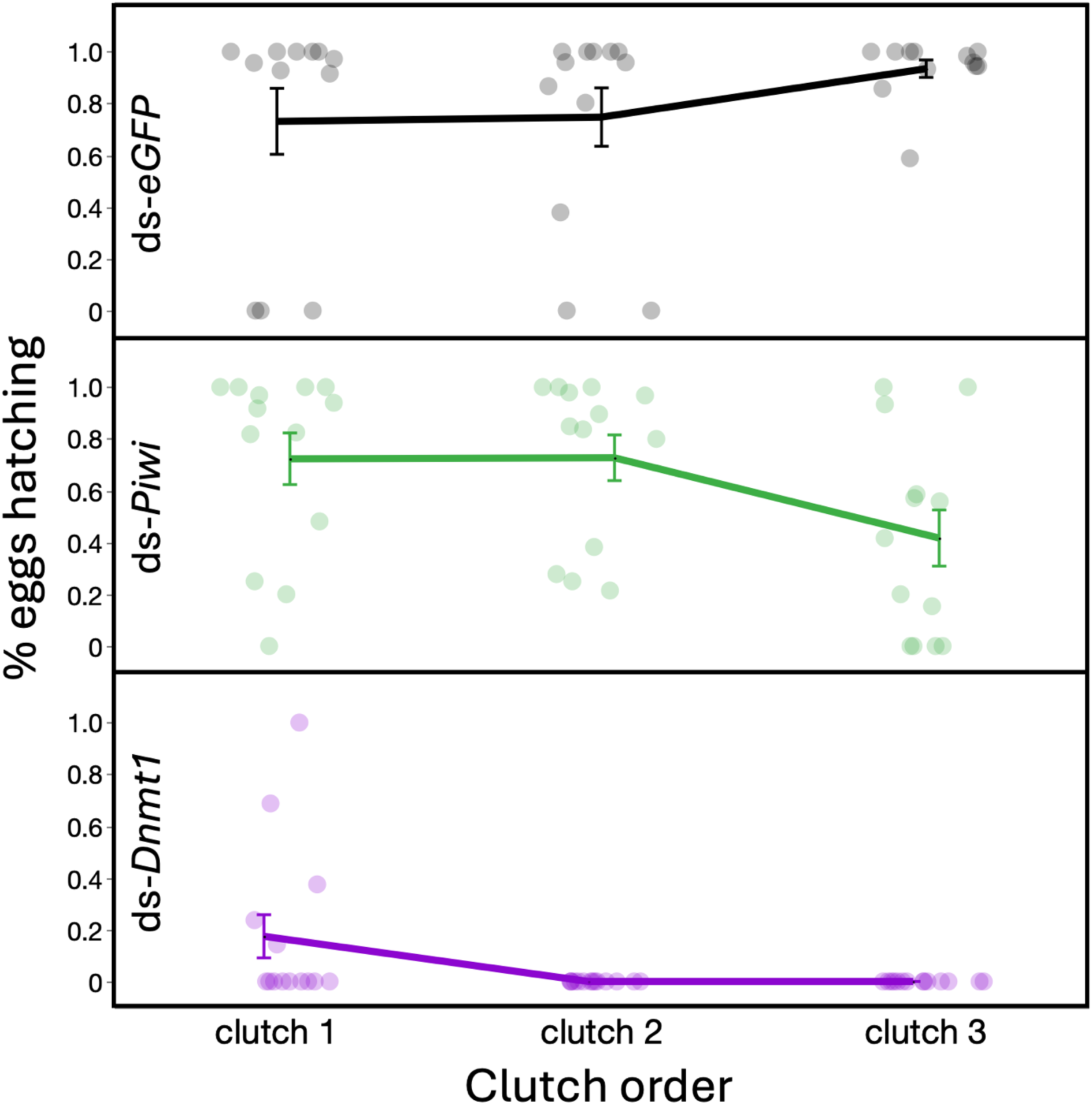
Both time and treatment influenced the hatch rate of eggs laid by females treated with dsRNA after adult emergence (repeated measures ANOVA_time*treatment_: p < 0.001). Eggs laid in the first three clutches by females treated with ds-*eGFP* (green rectangles) all had high percentages of eggs that hatched (n=12 females). Eggs laid in the first two clutches of eggs by females treated with ds-*Piwi* (blue diamonds) had hatch rates indistinguishable from the hatch rates of the control females. The hatch rate in the third clutch, however, was significantly reduced compared to controls (n = 13 females). Repeated measures ANOVA time*treatment p < 0.001. Eggs laid in the first clutch of eggs laid by females treated with ds-*Dnmt1* (red circles) had very low hatch rates and none of the eggs in the second and third clutches of eggs hatched (n = 14 females). Markers represent the mean values +/- standard error. Dots represent individual females.

### Dnmt1, Piwi, and Vasa are expressed in different populations of cells within the ovariole

We localized mRNA for our three target genes within the ovarioles of sexually mature females using HCR (Chen et al. 2018). We found that these three mRNAs were localized in overlapping but different populations of cells within the ovariole (Figure 2). *Dnmt1* was expressed at low levels in the trophocytes within the germarium at the tip of the ovariole (Figure 2B and 2C). More intense staining with the *Dnmt1* probes was observed in very small cells (Figure 2C, yellow arrows) in the region of the ovariole that contains the primary oocytes (Ewen-Campen et al. 2013) embedded within pre-follicular cells located at the base of the germarium. *Dnmt1* staining persisted in the maturing oocytes as they moved down the pedicel towards the oviduct. *Vasa* expression pattern largely overlapped with *Dnmt1*, with the signal observed in the trophocytes as well as the young and maturing oocytes (Figure 2B and 2D). The major difference we observed was that the smallest of the primary oocytes had little to no staining with the *Vasa* probe (Figure 2D yellow arrows). *Piwi* staining was unique in that signal from the *Piwi* probe was only observed within the trophocytes. There was reduced *Piwi* staining in the oogonia or oocytes. None of the three probes labeled the somatic follicular cells surrounding the oocytes. Broadly, the pattern of *Vasa*, *Piwi*, *Dnmt1*, and DAPI for ds*Vasa* samples is consistent with the control (ds-*eGFP*; Figure 2M, N, and P) reinforcing that *Vasa* gene expression reduction has little direct effect on oogenesis.

**Figure 2.**
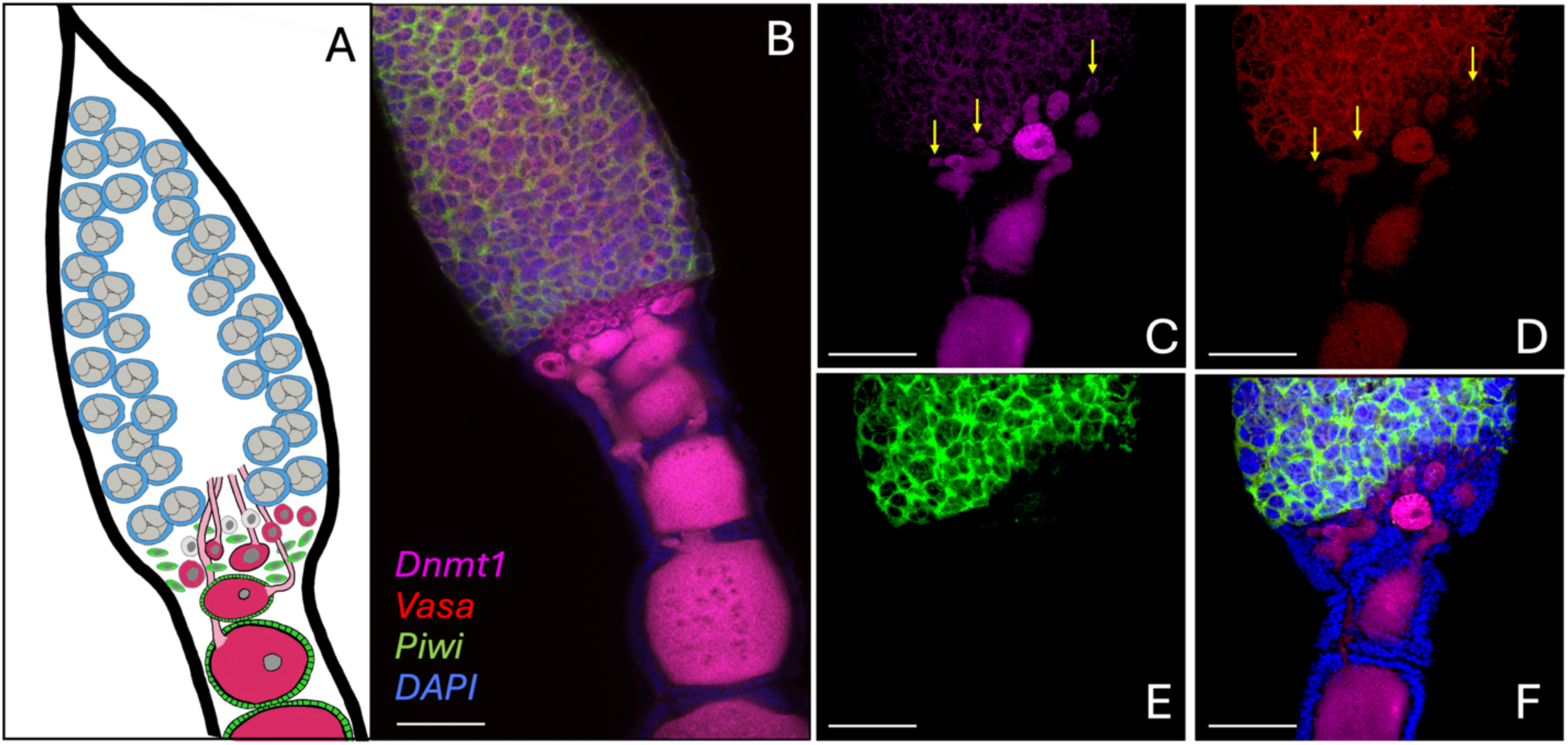
*Dnmt1*, *Vasa*, and *Piwi* were expressed in both overlapping and distinct cell types in the adult ovariole. (A) Schematic diagram of an *O. faciatus* ovariole. *Oncopletus faciatus* have telotrophic, meroistic ovaries containing seven ovarioles per ovary. During the fourth instar nymphal stage of development, oogonia set aside during embryogenesis divide mitotically to form nutritive cells, the trophocytes, and meiotically to form gametes, the oocytes. The trophocytes remain at the apical tip of the ovariole (blue cells with large gray nuclei). In the region below the trophocytes there is a region that contains the oogonia (small gray cells), and ooctyes (dark pink cells) that develop from small primary ooctyes to maturing ooctyes that move into the pedicel of the ovariole. Oocytes are connected to the trophocytes through trophic cords (light pink structures; Huebner 1981). The oogonia and ooctyes are embedded in pre-follicular cells (green cells) that envelope the maturing ooctyes as they descend and form a follicular epithelium. (B-F) HCR images of ovarioles from sexually mature stock females labeled with probes that recognize *Vasa* (red), *Piwi* (green), and *Dnmt1* (magenta) RNA. The ovarioles were also stained with DAPI (blue) to localize nuclei. (B) Composite image of control ovariole imaged at 20X. (C-F) Detailed image of the region containing the oogonia and primary oocytes within a control ovariole. (C) *Dnmt1* signal, (D) *Vasa* signal, (E) *Piwi* signal, and (F) composite image. 20X magnification. Scale bar = 100 mm.

### Dnmt1 and Piwi knockdown affected the different stages of oocyte development

*Dnmt1*, *Piwi*, and, *Vasa* RNAi knockdowns had variable effects on ovariole structure in females treated as fourth instar nymphs, during ovarian development. In our ds-*eGFP* control females, the structure and HCR staining pattern for all three genes was indistinguishable from untreated females (Figure 3A-D). As we have observed previously, ovarioles dissected from sexually mature females that developed following ds-*Dnmt1* treatment were very small with no maturing oocytes visible. The HCR staining of these ovarioles indicated a general breakdown in ovariole structure (Figure 3E-H). That is, the ovarioles looked as if all cell types are being affected rather than one specific cell type. As expected, there was no staining with the *Dnmt1* probe. Staining levels were also reduced and diffuse for the *Vasa* and *Piwi* probes. This was particularly notable in the trophocytes where there was reduction in the clear delineation between the multinucleate cells. In ds-*Piwi* treated females, the ovarioles were also smaller with few developing oocytes. Additionally, as was apparent from the *Dnmt1* and *Vasa* HCR staining in the ds-*Piwi* treated females (Figure 3I-L), the oocytes that were present were not maturing as expected. While the small primary oocytes were embedded in the pre-follicular cells as in the controls, growing oocytes did not enter into the stalk or become enveloped by follicular epithelium cells. In all samples, we observed the same pattern as illustrated here with the oocyte accumulating some yolk, but never successfully entering the ovariole stalk, growing larger, or taking on the typical shape of a maturing oocyte. As expected, the ovarioles from ds-*Vasa* treated females had a structure and staining patterns that did not differ from controls (Figure 3M-P).

**Figure 3.**
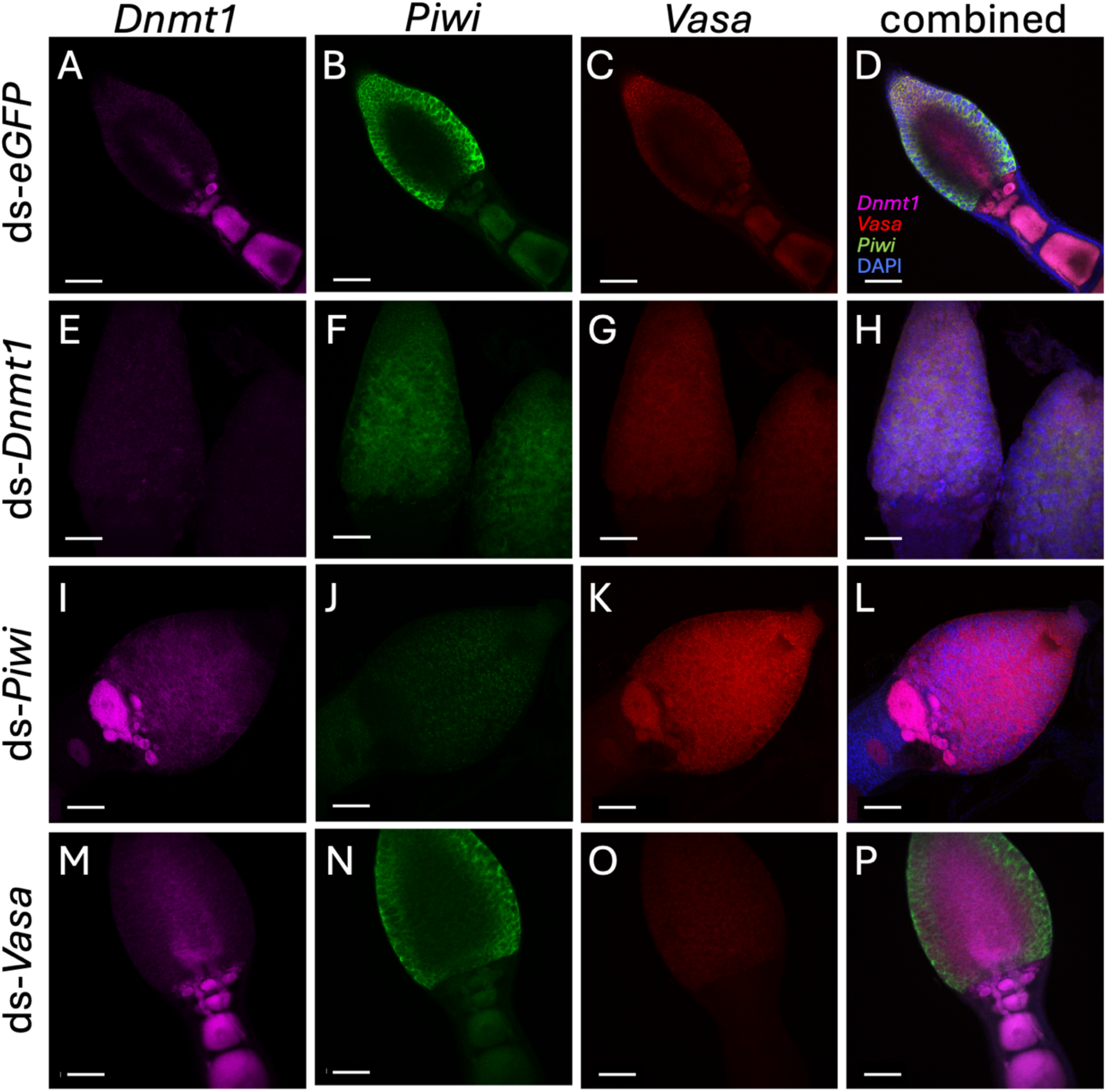
Gene knockdown of *Vasa*, *Piwi*, and *Dnmt1* influenced gene expression in both predictable and unexpected ways in ovaries. HCR staining of ovarioles dissected from sexually mature females treated with dsRNA during larval development. (A-D) Ovarioles from ds-*eGFP* treated females showed the same staining pattern and structure as the ovarioles from colony females. (A) *Dnmt1* mRNA (magenta) was localized to the oocytes, including the earliest stages, (B) *Piwi* mRNA (green) was confined to the trophocytes, and (C) *Vasa* mRNA (red) was localized to the trophocytes and maturing ooctyes. In every RNAi knockdown treatment, the corresponding signal is reduced compared to controls, indicating successful knockdown. (E-H) *Dnmt1* knockdown ovarioles looked disorganized and the signal from all the probes was less intense than the other treatments suggesting a knock-on effect on of this treatment. (I-L) *Piwi* knockdown females had oocytes that initiated development but failed to progress as normal and did not move into the stalk or become enveloped in the follicular epithelium. (M-P) *Vasa* knockdowns were not distinguishable from controls other than the reduction in signal from the Vasa probe. 20X magnification. Scale bars = 100 mm.

While knockdown of *Dnmt1* eliminates maturing oocytes, we wondered if reduction in *Dnmt1* expression during ovarian development also eliminated the earliest stages of oocytes which are embedded within the pre-follicular cells and difficult to distinguish. Unpublished work in our lab had suggested that anti-β-catenin antibody labeled developing oocytes, including at the earliest stages of development. This preliminary observation led us to examine the distribution of β-catenin in the ovaries of females treated with ds-*Dnmt1* or ds-*Piwi*. In ovarioles from ds-*eGFP* treated control females labeled with anti-β-catenin antibody, there was nuclear staining in a distinct population of small cells in the region below the trophocytes where pre-follicular cells and primary oocytes are located (Ewen-Campen et al. 2013) but staining was absent from the nuclei of surrounding pre-follicular cells (Figure 4A-C). Anti-β-catenin staining persisted into the nuclei of maturing oocytes. In ovarioles of sexually mature females treated with ds-*Dnmt1* or ds-*Piwi* as fourth instar nymphs, we found that there was no effect on the population of cells that are stained with the anti-β-catenin antibody (Figure 4D-I). In both treatments the population of anti-β-catenin positive nuclei was apparent embedded within the pre-follicular cells.

**Figure 4.**
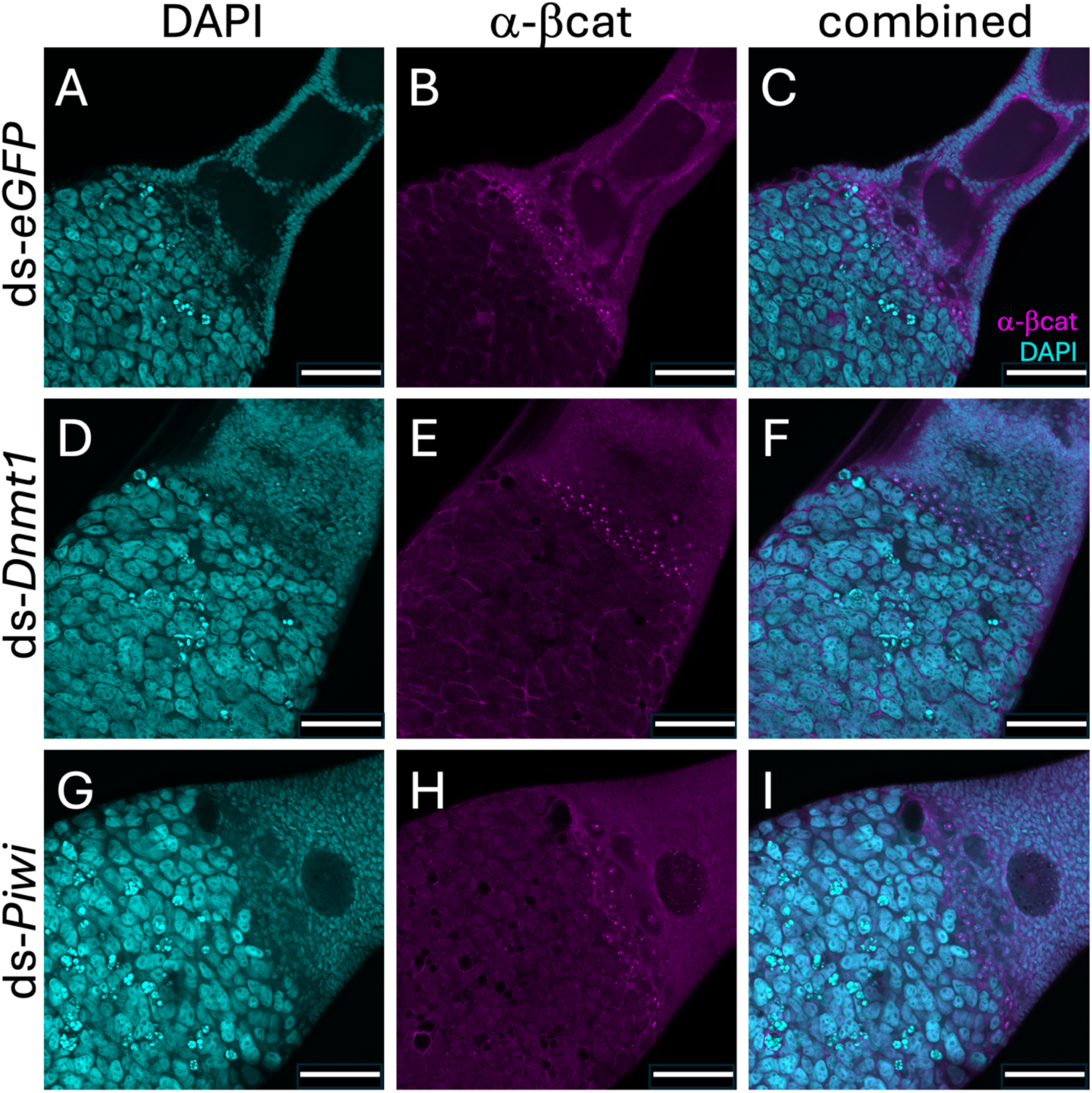
β-catenin antibody stains a population of cells, thought to be oocytes, that are embedded in pre-follicular cells in the germarium. Anti-β-catenin staining of ovariole tips from females knocked down during larval development. A-C. Ovarioles from sexually mature females treated with ds-*eGFP* as L4 larvae. The ovarioles looked typical, with developing oocytes moving through the pedicle. Anti-β-catenin antibody stained the nuclei of a population of cells that includes the oogonia and primary oocytes and the nuclei of developing oocytes. In ovarioles from sexually mature females treated with ds-*Dnmt1* (D-F) or ds-*Piwi* (G-I) as L4 larvae there were no developing oocytes within the pedicle, but the anti-β-catenin positive nuclei within the population of cells that includes the oogonia and primary oocyte remain with no detectable reduction in numbers. 20X magnification. Scale bar = 100 mm.

### All three target genes were required for male fertility

To test for the effect of *Vasa* and *Piwi* on spermatogenesis, we treated adult males with dsRNA and allowed them to mate with virgin females, using ds-*Dnmt1* and ds-*eGFP* as controls. Males molt into adults with functional sperm (Economopoulos and Gordon, 1971), so we mated males sequentially to observe the effect of our adult RNAi treatment as that would only affect newly developing sperm. We found that the fertility of males was negatively affected over time (Figure 5; repeated measures ANOVA, *F*_within_ = 17.365, d.f. = 3, p < 0.001), the change depended on treatment (F_between_ = 26.382, d.f. = 3, p < 0.001), and there was a significant interaction between treatment and time (F_time*treatment_ = 5.402, d.f. = 9, p < 0.001). As expected, based on previous research, *Dnmt1* knockdown had a severe effect on male fertility (Washington et al. 2021). The impact of *Piwi* knockdown was similar to that of *Dnmt1* knockdown, while *Vasa* knockdown appeared to be intermediate (Figure 5).

**Figure 5.**
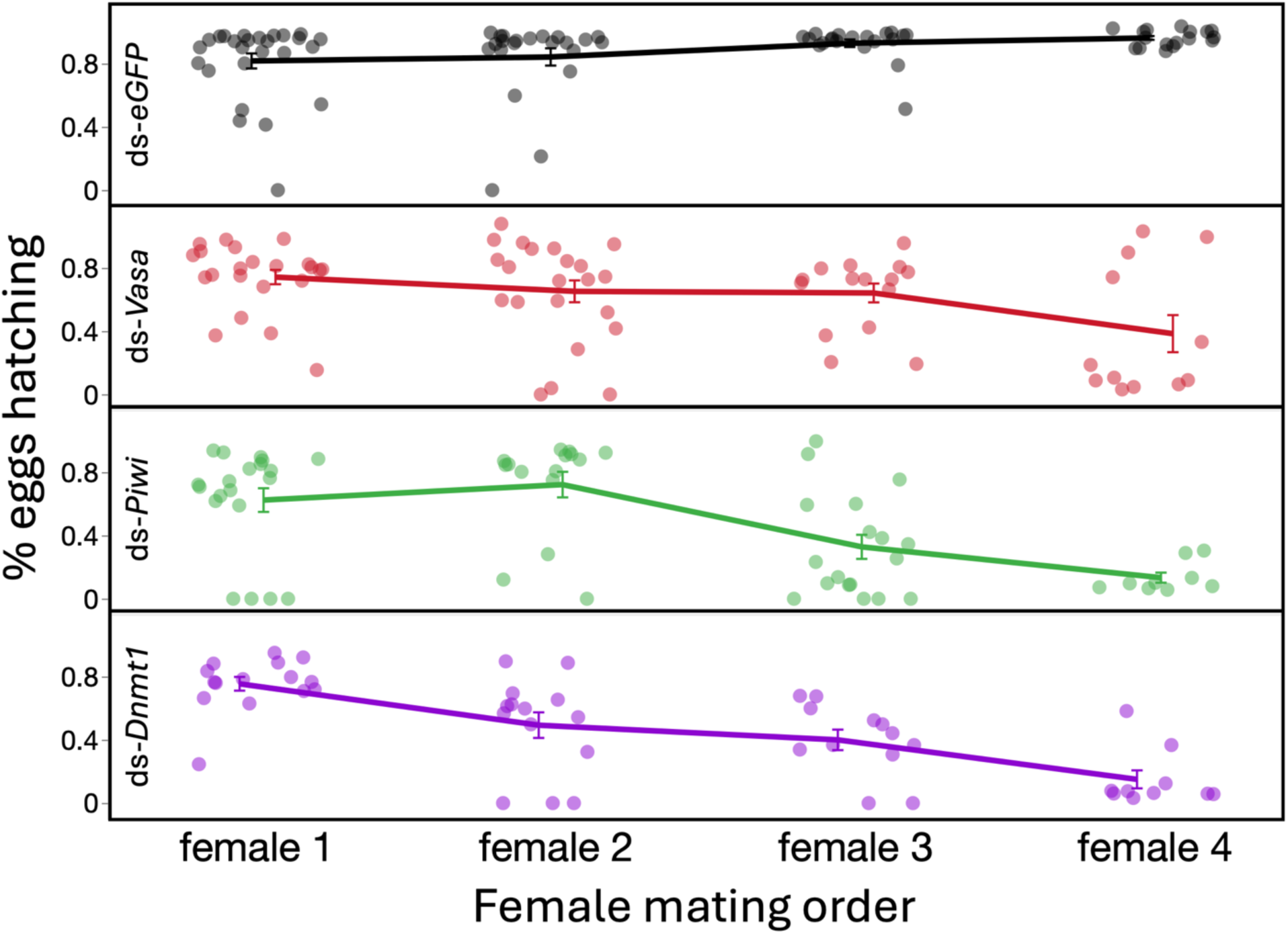
Both time and treatment reduced successful male mating. Male sperm supplies as measured by the percent of eggs laid that hatched decreased across time for all three dsRNA treatments (repeated measures ANOVA_time*treatment_ p < 0.001). The percent of fertilized eggs produced across the lifetime of females that mated to males across four weeks remained high for the ds-*eGFP* treated males (n = 26), but fell in females mated to RNAi treated males over multiple previous mates (ds-*Vasa* n = 23, ds-*Piwi* n = 22, ds-*Dnmt1* n = 16; repeated measures ANOVA_between_ p < 0.001.) Markers represent mean values. Dots represent individual males.

### Dnmt1, Piwi, and Vasa show distinct expression patterns in the testis tubule

In *O. fasciatus*, each testis tubule is organized from the earliest stages of spermatogenesis at the tip to the mature sperm at the base (Schmidt et al. 2001.) Thus, there exist bands of spermatocysts at particular developmental stages along the length of the testis tubule (Figure 6A, also see Ewen-Campen et al. 2013b Figure 6). We examined in which stages our target genes were expressed. There is a rosette of germline stem cells (GSCs) at the tip of the testis tubule that are connected to the stem cell niche within (Schmidt et al. 2001, 2002). The GSCs divide with one daughter cell leaving the rosette to become primary spermatogonia that are enclosed by a cyst cell. One of our most interesting results is that *Piwi* mRNA is concentrated in the rosette of GSCs (Figure 6B, E, G). After six mitotic divisions, the spermatogonia divide to form primary spermatocytes. The spermatocytes then divide meiotically to form haploid spermatids. *Dnmt1* mRNA signal was concentrated in the secondary spermatogonia (Figure 6B, C, and G) while *Piwi* and *Vasa* signal was most highly concentrated in the primary spermatocytes, with *Vasa* signal persisting in the secondary spermatocytes (Figure 6B, D, E, and G).

**Figure 6.**
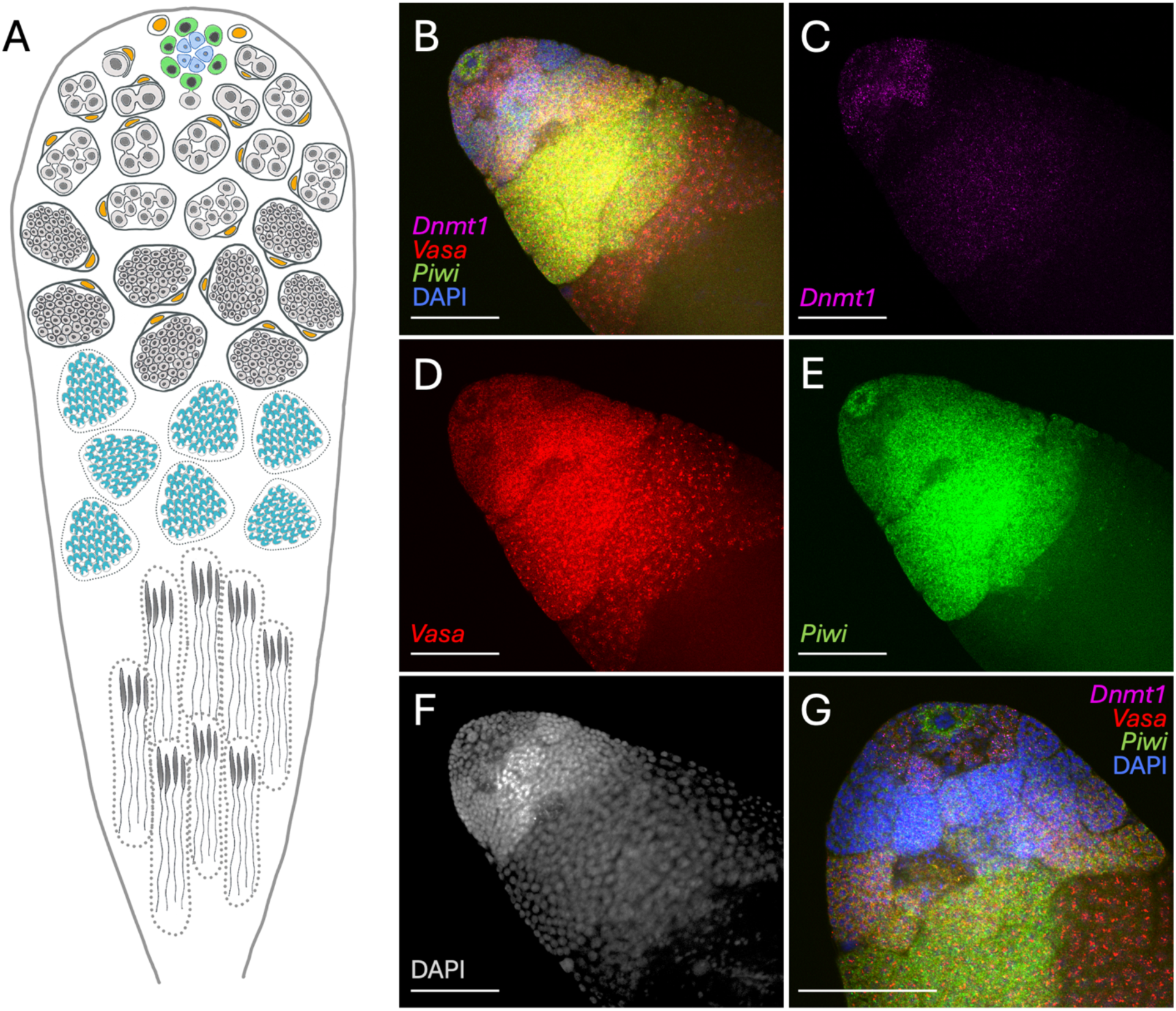
*Vasa*, *Piwi*, and *Dnmt1* were expressed in both overlapping and distinct cell types in *O. fasciatus* testes. (A) A schematic diagram of the stages of spermatogenesis in an *Oncopeltus* testis tubule (taken from Washington et al. 2021). In *O. fasciatus*, spermatogenesis proceeds from the tip of each of seven testis tubules within each testis. At the apical tip of a testis tubule there is a rosette of germline stem cells (green cells) surrounding a hub of niche cells (light blue cells, Schmidt et al. 2002). As spermatogonia (light gray) arise from division of the germline stem cells, they are enclosed by cyst cells (yellow). The spermatogonia undergo 6 mitotic transit amplification divisions. At the 64-cell stage the spermatogonia divide to form primary spermatocytes (dark gray; Economopoulos & Gordon 1971, Ewen-Campen et al. 2013). *Oncopeltus fasciatus* undergoes inverted meiosis (Viera et al. 2009). Primary spermatocytes undergo the first meiotic division to produce diploid secondary spermatocytes. The second meiotic division produce the haploid spermatids (turquoise cells) that then differentiate into spermatozoa. Identification of the stage of spermatogenesis can be accomplished by the number of nuclei in a cyst and nuclear morphology (F; Ewen-Campen et al. 2013). (B-G) HCR staining of sexually mature testis tubule from untreated males. The signal for the *Piwi* mRNA (green) was strong in the germline stem cell rosette (B, E), but *Piwi* mRNA was absent from the niche cells. In the region of the testis tubule where primary spermatogonia are undergoing the first transit amplification divisions there was little staining with any of the probes. Staining for *Dnmt1* (magenta) increased within the secondary spermatogonial cysts as they approach the 64 nuclei stage at which they will transition from spermatogonia to spermatocytes (C). While there was still some *Dnmt1* positive signal in the developing spermatocyte cysts, the signal is lower and the nature of the staining changes from punctate to more diffuse. *Vasa* (red) expression came on in the primary and secondary spermatocytes (D) and a positive signal was still evident in the developing spermatids. *Piwi* expression, which was absent in the early spermatogonial cysts, increased strongly in the developing spermatocyte cysts and ended abruptly in the region where the spermatocytes transition to spermatids (E). 20X magnification. Blue stain is DAPI. Scale bars = 100 mm.

When we examined the expression patterns of the three genes within the testis tubules of males that had been treated with ds-RNA after adult molt and then allowed to mate for 10 days, the phenotype was similar among the different treatments. In all knockdowns the testis tubules contained fewer developing spermatocysts, particularly at the apical end (Figure 7), but relative location of expression for the three genes was mostly maintained. Interestingly, in the *Dnmt1* knockdown males, the strong expression of *Piwi* in the GSC rosette was not apparent. While we examined multiple testis tubules (Supplementary Materials Figure S3), it would be difficult to make a conclusive statement that the rosette was absent without further study. The nuclei are so concentrated in this region of the testis tubule that it was typically very difficult to pick out the rosette nuclei above background staining in whole mounts stained with DAPI. However, this observation received support from the qRT-PCR results in which testis tubules from ds-*Dnmt1* treated males had significantly reduced expression of *Piwi* as well as reduced expression of *Dnmt1* (Supplementary Materials Figure S2).

**Figure 7.**
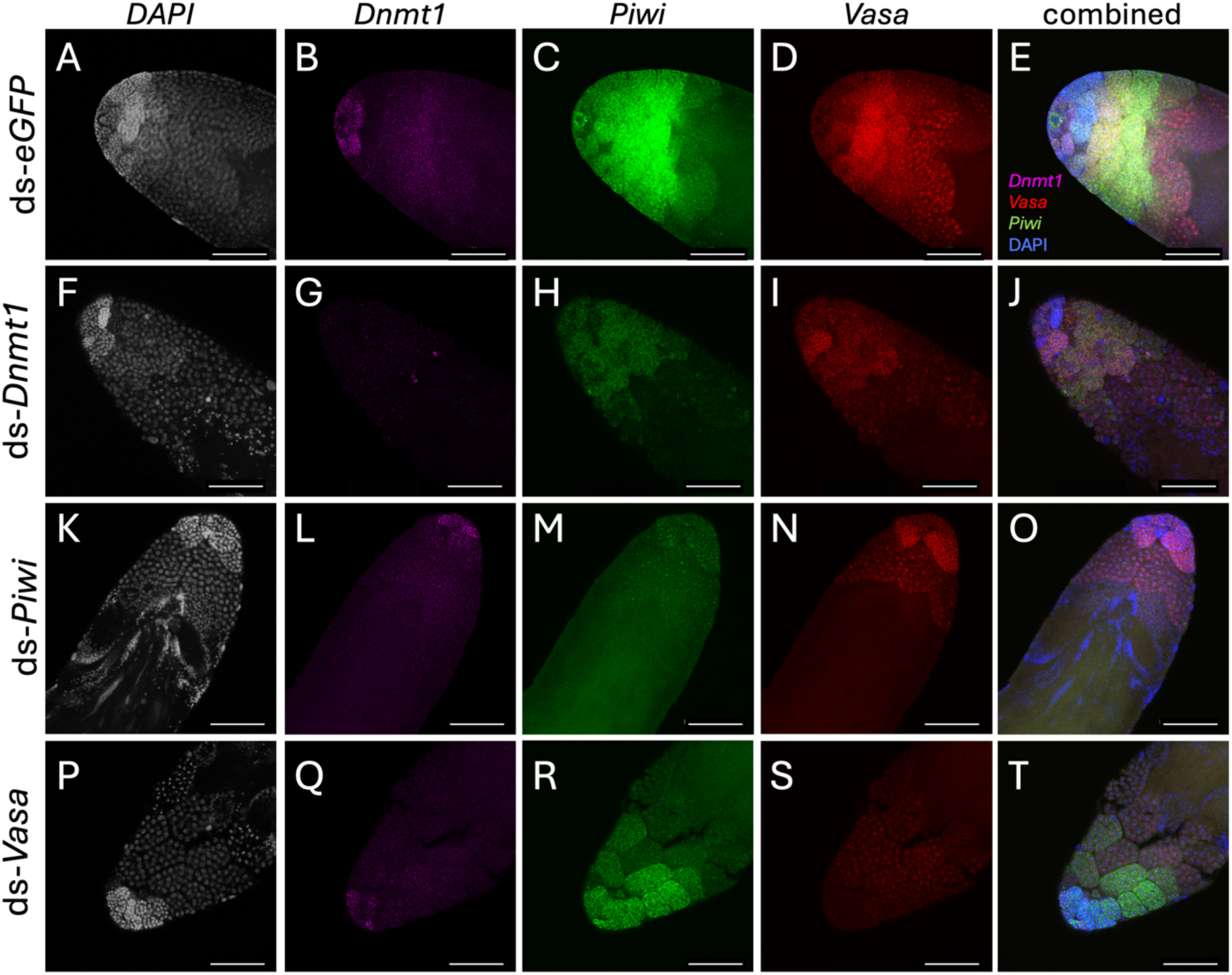
Gene knockdown of *Vasa*, *Piwi*, and *Dnmt1* influenced gene expression in a predictable way in testes. HCR of testis tubules from males treated with ds-RNA during sexual maturation. Males were then allowed to mate to investigate whether any or all of the genes were functional required to replenish sperm stores. (A-E) Testis tubule from a *ds-eGFP* treated male showed the typical pattern we have observed for gene expression in control testis tubules. *Dnmt1* (B; magenta) is expressed primarily in the secondary spermatogonia. *Piwi* (C; green) is expressed in the germline stem cell rosette at the apical tip and in the primary spermatocytes. *Vasa* (D; red) is expressed in the primary and secondary spermatocytes. In the *dsDnmt1* treated testis tubules (F-J), the testis tubules appeared to have fewer spermatocysts, but the pattern of expression of *Piwi*, *Vasa*, and *Dnmt1* was similar to controls. In both the *ds-Piwi* (K-O) and the *ds-Vasa* (P-T) knockdown testis tubules, the result was similar to the *dsDnmt1* with testis tubules that appeared to have fewer spermatocysts but no major change to the expression pattern outside of the target gene. While the organization of the expression pattern was comparable to the control testis tubules, the number of spermatocysts expressing each gene, particularly the Vasa positive spermatocysts, was observably reduced. 20X magnification. Blue stain is DAPI. Scale bars = 100 mm.

## Discussion

Given the fundamental role of germ cells in fitness, there exists a highly conserved core of genes across diverse groups of organisms, suggesting that this gene set is under high levels of evolutionary constraint (Fierro-Constaín et al. 2017, Piccinni & Milani 2023). However, when these genes are examined across the Metazoa beyond presence/absence, variation begins to emerge in the details of the biochemical processes by which these core genes regulate their definitive phenotype. Thus, to truly understand the evolutionary history and variation of these genes, it is critical to identify the specific cell types expressing these genes and functionally characterize their role between the sexes and across a diverse set of species. Here we focused on three genes within this set of germline core genes, *Vasa*, *Piwi*, and *Dnmt1* in both male and female *O. fasciatus*. We focused on the function of these genes in the development of gametes following establishment of the germline by localizing their mRNA and through gene expression knockdown using RNAi to extend our understanding of the specific gametogenic processes they are involved with and the specific cell types in which they are present. *Vasa* was present in germline cells. It was not required for oogenesis but was required during spermatogenesis. *Piwi* was present in germline cells and required for both oogenesis and spermatogenesis. *Dnmt1* was present in germline cells and required for both oogenesis and spermatogenesis. *Piwi* is expressed in the testis germline stem cells (GSCs), but both *Vasa* and *Dnmt1* expression was absent from these cells. These results, and those found for these genes in other organisms, highlight the unexpected variation found in the genetic networks required for gametogenesis and hint at a more nuanced evolutionary story underpinning the shared conserved gametogenic gene set in Metazoa.

Given its ubiquity of expression within germline cells across the Metazoa (Adashev et al. 2023), it is perhaps surprising that *Vasa* does not play a role in oogenesis in *O. fasciatus* (Ewen Campen et al. 2013b). However, the requirement of *Vasa* for oogenesis is variable across the insects (Adashev et al. 2023). *Vasa* is not required for oogenesis in the cricket *G. bimaculatus* (Ewen-Campen 2013a). A caveat for the functional analysis of *Vasa* in crickets and *O. fasciatus*, as well as the study presented here, however, is that knockdown occurred late in female germline development after primary oocytes are already present (Wick and Bonhag, 1955) This differs from *D. melanogaster* that have a female germline stem cell population in adults (Spradling et al. 2011). *Vasa* is required for GSC maintenance in *D. melanogaster*(Adashev et al. 2023), but an adult population of GSCs has never been described in *O. fasciatus* (Wick and Bonhag 1955, Ewen-Campen et al. 2013b.) Further studies in which *Vasa* is knocked down in female nymphs prior to the formation of primary oocytes would be required to determine if the function of *Vasa* in maintenance of GSCs in females is conserved. In this study we also confirmed the observation that *Vasa* is required for spermatogenesis in *O. fasciatus* and confirm that *Vasa* expression is highest in the secondary spermatogonia (Ewen-Campen et al. 2013b). *Vasa* mRNA is also detectible in primary and secondary spermatocytes, although low levels of expression were seen in earlier stages of development. Ewen-Campen et al. (2013b) suggest a role for *Vasa* in controlling meiosis, as is seen in mice (Tanaka et al. 2000), humans (Medrano et al. 2012), and *D. melanogaster* (Davis 1971). In our study, we saw a general reduction in the number of developing spermatocysts following treatment with *Vasa* dsRNA but we did not see anything that speaks definitively to a mechanism by which *Vasa* is working to reduce spermatogenesis.

*Piwi* was required for both oogenesis and spermatogenesis in *O. fasciatus*. *Piwi* is required for the self-renewal of GSC in *D. melanogaster* (Juliano et al. 2011, Santos et al. 2023). Given that *O. fasciatus* females do not have GSCs, we predicted that *Piwi* would not be required for oogenesis. However, *Piwi* knockdown did result in reduced fertility in females. This was not related to a reduction the number of primary oocytes. *Piwi* mRNA expressed strongly in the trophocytes, with low levels of signal in the youngest oocytes. The mechanism by which *Piwi* allows proper progression of the oocyte through the ovariole remains unknown. The effect of *Piwi* on hatch rates demonstrates that Piwi knockdown does not affect embryo development. Eggs that received *Piwi* dsRNA after a critical period progressed through oocyte maturation and resulted in viable embryos. The first two clutches of eggs represented oocytes that were in the later stages of maturation at the time of treatment. The drop off in hatch rate in the ds*Piwi* treated females suggests a requirement for *Piwi* during the earliest stages of maturation. One of our most intriguing observations in our *Piwi* knockdown females was vitellogenic eggs that never entered the pedicel and did not appear to associate properly with the follicular epithelial cells, leading to speculation that *Piwi* knockdown interfered with communication between the germ and soma. The impact of *Piwi* knockdown on male fertility likely arose due to the conserved function of *Piwi* on GSCs. In *O. fasciatus* there is a rosette of cells consisting of niche cells surrounded by GSCs at the tip of the testis tubule (Schmidt et al. 2001, 2002). In our images, a layer of *Piwi* expressing cells surrounding a group of interior unlabeled cells presented exactly as expected for the GSCs. As in all our knockdown testes, there was a reduction in the number of developing spermatocysts that we predicted was due to loss of the GSCs but no obvious defect at later stages of spermatogenesis. Further research will elucidate the mechanism by which *Piwi* controls the progression of gametes through both the ovariole and testis tubule of *O. fasciatus*.

Our *Dnmt1* results here both agreed with our previous studies and produced two novel observations. First, as with previous experiments, we observed no oocytes when *Dnmt1* expression is reduced in fourth instar nymphal females. However, in previous studies, we were unable to determine if the primary oocytes were absent or not developing past the primary oocyte stage. Here, β-catenin was localized in the nuclei of a population of cells embedded in the prefollicular cells at the base of the germarium. The nuclear localization persists as oocytes develop, leading to the inference that the cells with β-catenin nuclear localization are oocytes, although not directly demonstrated in this study. The persistence of this population of cells in the *Dnmt1* knockdown females indicates that *Dnmt1* functions in oocyte maturation and is required for oocytes to move from the primary oocyte stage to the vitellogenic stage of development. As with *Piwi*, it may be that *Dnmt1* is also required for the formation of the primary oocytes in the nymphal ovary but we did not perform the knockdown at an early enough stage of development to observe that function. The second novel observation in this study was the loss of *Piwi* staining in the GSC at the tip of testis tubule following knockdown of *Dnmt1* expression. Our imaging did not allow us to separate if *Dnmt1* is required for *Piwi* expression, or if *Piwi* expression is reduced because *Dnmt1* plays a direct role in the maintenance of the GSCs or the niche.

The interaction between *Piwi* and *Dnmt1*, as well as the similar patterns of conservation across the insects (Bewick et al. 2017, Santos et al. 2023), is interesting because both are involved in genome defense and fertility and thus have critical fitness functions. This appears to be contradictory to the evolutionary patterns in which we see variation, including loss of function of the gene product in gametogenesis.

Our paper highlights the need for integrated studies of expression and function when examining genes that are consistently present and therefore considered to be highly conserved. Moreover, we need more studies on organisms with different reproductive ecologies and physiologies to truly understand evolutionary patterns and processes around these genes. Our research suggests that plasticity in development, with the same gene sequences showing variability in expression or function, may be more common than previously supposed which will only be revealed with studies of diverse organisms. For example, *D. melanogaster* is one of the only insects in which ovarian GSCs have been documented (Spradling et al. 2011). Why are female GSCs so unusual? Spradling et al. 2011 suggest that this may be an evolutionary response to unpredictable and ephemeral resources for reproduction. It may also be that other species have female GSCs but they have not been identified yet. We posit that a requirement for *Vasa* in oogenesis post adult maturation is a clue to which species have GSCs.

We chose to study gametogenesis because it is predicted to be resistant to alteration given its relationship to fitness. Yet whilst it is logical to predict conservation of core genes involved in gametogenesis, it is also clear that variation in the timing and biochemical partners through which these genes act is common. For example, *Vasa* is largely conserved at a phenotypic level. It is required for germ cell fate in most organisms (Ewen-Campen et al. 2010, Chen et al. 2025). But the biochemical pathway by which *Vasa* acts can be variable, as demonstrated by the *Oskar*-independent mechanism proposed in the pea aphid (Lin and Chang 2025, Kao et al. 2026). This is in some ways surprising, given that we would predict that any mutation, such as a change that affected the network through which *Vasa* specifies germ cell fate, that interfered with *Vasa*’s function in gametogenesis would be strongly selected against and eliminated from the population. *Dnmt1* poses a different case. The biochemical action of *Dnmt1*, the maintenance of CpG DNA methylation marks following semi-conservative replication, is highly conserved across taxa (Schmitz et al. 2019), including insects. However, the phenotypic function in gametogenesis has been evolutionarily uncoupled from this enzymatic activity (Schulz et al. 2018, Bewick et al. 2019, Amukamara et al. 2020, Washington et al. 2021). Further details into the mechanism by which DNMT1 controls oogenesis and gametogenesis is required to fully understand the selective pressures leading to the role of DNMT1 in reproduction within the insects. As is apparent from these examples, conservation at one level does not imply conservation at another. The importance of this variation for evolution is unclear and deserves further study.

## Acknowledgements

The authors would like to thank Dr. Elizabeth Duncan for her sharing her protocols and expertise on HCR staining of insect gonads. We also thank Muthugapatti Kandasamy at the University of Georgia Biomedical Microscopy Core for help with the confocal microscopy.

## Competing Interests

No competing interests declared.

## Funding

This work was funded by the USDA Agricultural Research Service Non-Assistance Cooperative Agreement #58-6080-9-006 “Managing whiteflies and whitefly-transmitted viruses in vegetable crops in the southeastern U.S.” Any opinions, findings, conclusions, or recommendations expressed in this publication are those of the author(s) and do not necessarily reflect the view of the U.S. Department of Agriculture.

## Data and resource availability

The datasets analyzed and the original image files for this study will be made available by the authors through the publicly available Dryad Digital Repository (hyperlink to be added)

## Notes

### Competing Interest Statement

The authors have declared no competing interest.

