## supplemental materials for "Conserved genes with variable expression and function: *Vasa*, *Piwi*, and *Dnmt1* in *Oncopeltus fasciatus* gametogenesis"

Supplementary Materials

Table S1. Primer sequences used to generate RNAi for injection and to quantify expression levels using quantitative Real Time PCR

Primers to produce PCR products for transcription reaction to produce ds-RNA. T7 promoter sequences required for transcription were included on the gene-specific primer sequences as described for the *Ambion MEGAscript kit*.

| Gene | Sense primer | Anti-sense primer |
|---|---|---|
| *Dnmt1* | TGATGCTCGGCCTCAAAACAAGAT | ACTCCAGGAGGTGGAACAGTAGTCT |
| *Piwi* | ATGGGACCAAATAAGGAACACGATCCGT | TTGTGAGCATACTGTACTTGAGCAGGA |
| *Vasa* | AGGACTGGCAATGATGGTAGA | GTGCAAATTTCCTGGCTTCAT |

Primers for qRT-PCR

| *Gene* | Sense primer | Anti-sense primer |
|---|---|---|
| *Dnmt1* | GCTTGGACAAAGGCTACTACT | CTTCGTGGTCCCTTATCCTTATC |
| *Piwi* | GGAAGAGGTTCCGAGATGATTG | ACTCGGTATGGCTTGTTGTTAT |
| *Vasa* | CTGTTGCTCCTCAGGTTATT | CATTAAGCCTTCCAGGAGTAG |
| *Actin* | CTGTCTCCCGAAAGAGAATATG | TCTGTATGGATTGGAGGATCTA |

Figure S1.

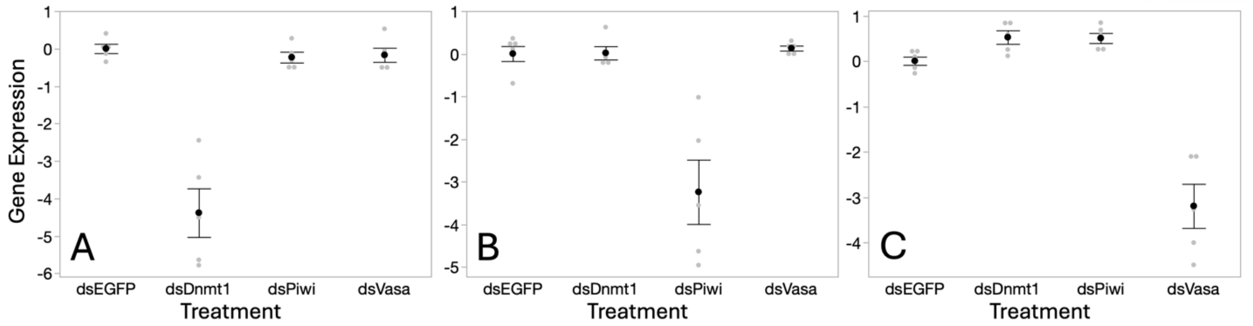


Figure S1. We tested expression levels of all three target genes in the ovaries of sexually adult females that had been treated with dsRNA as L4 nymphs. In each case, the gene targeted with the dsRNA had reduced expression. When there was a significant overall effect of treatment, we compared means from individual treatments to the control group of ds*eGFP* using Dunnett's Method. (A) *Dnmt1* expression was significantly affected by treatment ($F = 36.703$, d.f. = 3, 16, $p < 0.001$). *Dnmt1* expression was reduced in the ds*Dnmt1* treated females compared to control treatment ($p < 0.001$), but ds*Piwi* ($p = 0.936$) and ds*Vasa* ($p = 0.972$) treatment did not affect *Dnmt1* expression. (B) *Piwi* expression was affected by treatment ($F = 17.312$, d.f. = 3, 16, $p < 0.001$) and was significantly reduced in the ds*Piwi* treated females compared to controls ($p < 0.001$) but neither of the other treatments affected *Piwi* expression (ds*Dnmt1* $p = 1.00$; ds*Vasa* $p = 0.991$). (C) *Vasa* expression was affected by treatment ($F = 45.694$, d.f. = 3, 16, $p < 0.001$). *Vasa* expression was significantly reduced in the ds*Vasa* treated females compared to controls ($p < 0.001$), but not in ds*Dnmt1* ($p = 0.390$) or ds*Piwi* ($p = 0.420$) treated females.

Figure S2.

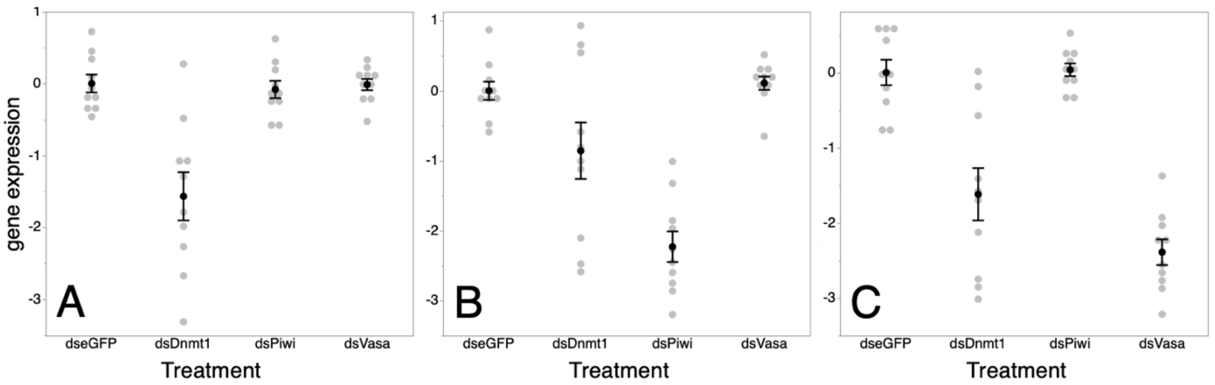


Figure S2. We tested expression levels of all three target genes in the testes of sexually adult males that had been treated with dsRNA during sexual maturation and then allowed to mate for 10 days following treatment. In each case, the gene targeted with the dsRNA had reduced expression. When there was a significant overall effect of treatment, we compared means from individual treatments to the control group of ds*eGFP* using Dunnett's Method. (A) *Dnmt1* expression was significantly reduced in testes from males treated with ds*Dnmt1*. The overall ANOVA was significant ($F = 15.991$, df = 3,36, $p < 0.001$). When we compared means from individual treatments to the control group, the ds*Dnmt1* treated males were significantly different ($p < 0.001$) from control males. But expression levels in testes from males treated with either dsPiwi ($p = 0.981$) or dsVasa ($p = 1.000$) were not different from control males. (B) *Piwi* expression was significantly affected by treatment ($F = 19.687$, df = 3,36, $p < 0.001$). As expected, the testes from males treated with ds*Piwi* had significantly reduced expression compared to the control group of ds*eGFP* ($p < 0.001$). The expression of *Piwi* was also reduced in testes from males treated with ds*Dnmt1* ($p = 0.046$). *Piwi* expression was not affected by treatment with ds*Vasa* ($p = 0.978$.) (C) Similarly to what was observed with *Piwi* expression, *Vasa* expression was significantly affected by treatment ($F = 31.434$, df = 3,36, $p < 0.001$). As expected, the testes from males treated with ds*Vasa* had significantly reduced expression of Vasa compared to the control group of dseGFP ($p < 0.001$). The expression of *Vasa* was also reduced in testes from males treated with ds*Dnmt1*. *Vasa* expression was not affected by treatment with ds*Piwi* ($p = 0.999$.)

Figure S3.

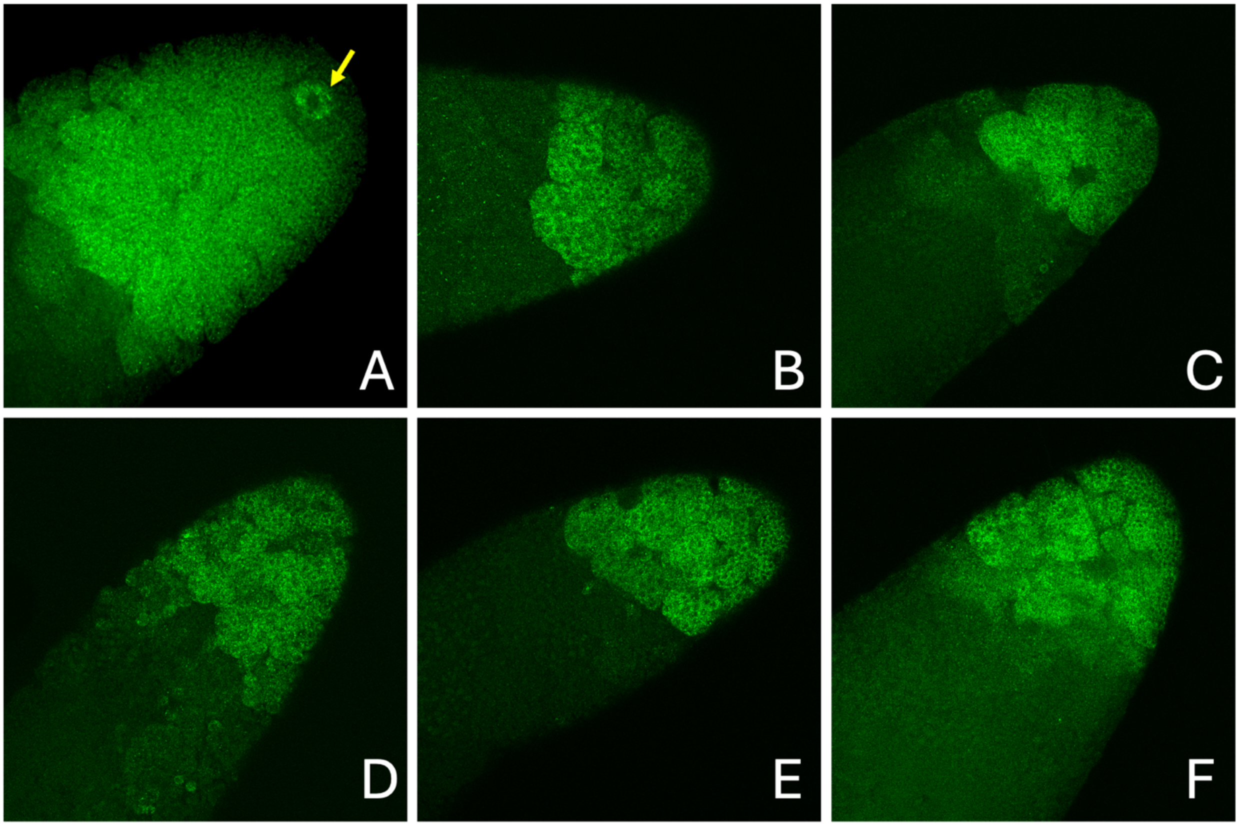


Figure S3. A selection of HCR images of testis tubules from adult males treated with ds*eGFP* (A) or ds*Dnmt1* (B-F) showing the probe for *Piwi* mRNA. The rosettes staining that is typically seen at the tip of the testis tubule of control males (A, arrow) is absent from the ds*Dnmt1* treated testis tubules, even though the developing spermatocysts did have a positive signal with the *Piwi* HCR probe (B-F). All images 20X magnification.
